# Phage activity against *Staphylococcus aureus* is impaired in plasma and synovial fluid

**DOI:** 10.1101/2023.05.11.540358

**Authors:** Michele Mutti, David Sáez Moreno, Marcela Restrepo-Córdoba, Zehra Visram, Grégory Resch, Lorenzo Corsini

## Abstract

S. aureus is a pathogen that frequently causes severe morbidity and phage therapy is being discussed as an alternative to antibiotics for the treatment of S. aureus infections. In this in vitro and animal study, we demonstrated that the activity of anti-staphylococcal phages is severely impaired in plasma and synovial fluid. Despite phage replication in these matrices, lysis of the bacteria was slower than phage propagation, and no reduction of the bacterial population was observed. This phage inhibition is due to a 99% reduction of phage adsorption, already at 10% plasma concentration. Coagulation factors bind S. aureus resulting in the formation of aggregates and blood clots that protect the bacterium from the phages. This was confirmed by the finding that purified fibrinogen is sufficient to impair phage activity. In contrast, dissolution of the clots by tissue plasminogen activator (tPA) partially restored phage activity. Consistent with these in vitro findings, phage treatment did not reduce bacterial burdens in a neutropenic mouse S. aureus thigh infection model. In summary, phage treatment of S. aureus infections may be fundamentally challenging, and more investigation is needed prior to proceeding to in-human trials.

## 1. Introduction

*Staphylococcus aureus* (*S. aureus*) causes systemic infections ^[1]^, ulcers ^[2]^, atopic dermatitis ^[3]^ and implant associated infections ^[4]^. Phage therapy, i.e the therapeutic use of species-specific bacterial viruses called bacteriophages (phages), has been proposed for the treatment of *S. aureus* infections, mainly because Staphylococcal phages often have a wide host-range, are active against biofilms in vitro and can evolve towards increased virulence ^[5]–[9]^.

These *in vitro* data demonstrate potential of *S. aureus* phage therapy, and some in-human studies have reported promising outcomes ^[10]^. Several in vivo studies of staphylococcal phages suggest that their lytic activity might be reduced in presence of plasma/serum or synovial fluid. To our knowledge, the first reference to the inhibitory effect of serum on phage lysis was made from in vitro observation by ^[11]^ and confirmed again very recently for *S. aureus* ^[12],[13]^. Inhibition of phage activity on *S. aureus* in vivo was reported in a mouse bacteremia model ^[14]^, where mice survived *S. aureus* challenge if the treatment was administered simultaneously, but not if the treatment was administered 30 min post challenge. This indicates that the adsorption of phages to *S. aureus* may be severely reduced in plasma. ^[12],[13]^ However, to our knowledge, characterization of the inhibition of *S. aureus* phage activity in biofilm-like aggregates formed in plasma or synovial fluid has not yet been reported.

*S. aureus* has several virulence factors that use the human coagulation system to its advantage, by either inducing coagulation surrounding the bacteria (*via* coagulase A), or by binding fibrin and fibrinogen a process called clumping (*via* clumping factor A and B, ClfA and ClfB, respectively). This greatly influences both the colonization of the site of infection and the evasion of the immune response ^[15]^. In the context of periprosthetic joint infections (PJI), *S. aureus* also forms aggregates in synovial fluid ^[16],[17]^. This may enhance the stability of S. aureus biofilms ^[18]^, its capacity to evade the immune system ^[19]^ and its tolerance to antibiotics ^[16],[18]^.

In this study, *we* tested wild type phages and phages evolved for increased virulence ^[9]^ *in vitro* in plasma and synovial fluid, and found that phage lytic activity is inhibited in these conditions. We show that masking of the phage receptor(s) on the surface of *S. aureus* cells by fibrinogen inhibits phage absorption. Furthermore, the thrombolytic drug tissue plasminogen activator (tPA) was found able to restore phage adsorption and lytic activity *in vitro*. These findings are consistent with the lack of efficacy of high-dose of *S. aureus* phages in an in vivo mouse thigh infection model. Overall, while phage therapy to treat *S. aureus* infections may have some potential based on in vitro results, plasma inhibition represents a limitation to their systemic efficacy *in vivo* and needs to be investigated further.

## 2. Results

### 2.1. Inhibition of phage activity in human and guinea pig plasma

To better understand the extent of phage inhibition in plasma, we measured the antibacterial activity of phages in broth supplemented with human plasma. Multiple, previously described individual phages and cocktails were used, including the Podoviridae phage P66, *two Herelleviridae* phages (PM4 and PM93), as well as two phage cocktails (cPM398, composed of P66 and vB_SauH_2002, and cPM399, composed of PM4, PM93 and PM56), in a bacterial growth inhibition assay in brain-heart infusion (BHI) medium supplemented with increasing amounts of human plasma.

All tested phages and cocktails showed impairment of their lytic activity, starting already at 0.5% plasma (Fig. 1a). While staphylococcal growth inhibition was observed over 48h in growth medium without plasma, bacterial regrowth was observed in presence of 0.5% plasma. PM93 was less affected than other phages, as bacterial growth over 48h was less pronounced at 0.5% plasma than for the other phages. The bacterial growth kinetics in presence of P66 were also very similar in absence and presence of 0.5% plasma, (initial lag phase before outgrowth longer) than for reactions containing 5% or 10% plasma, where P66 appeared completely inactive.

**Figure 1.**
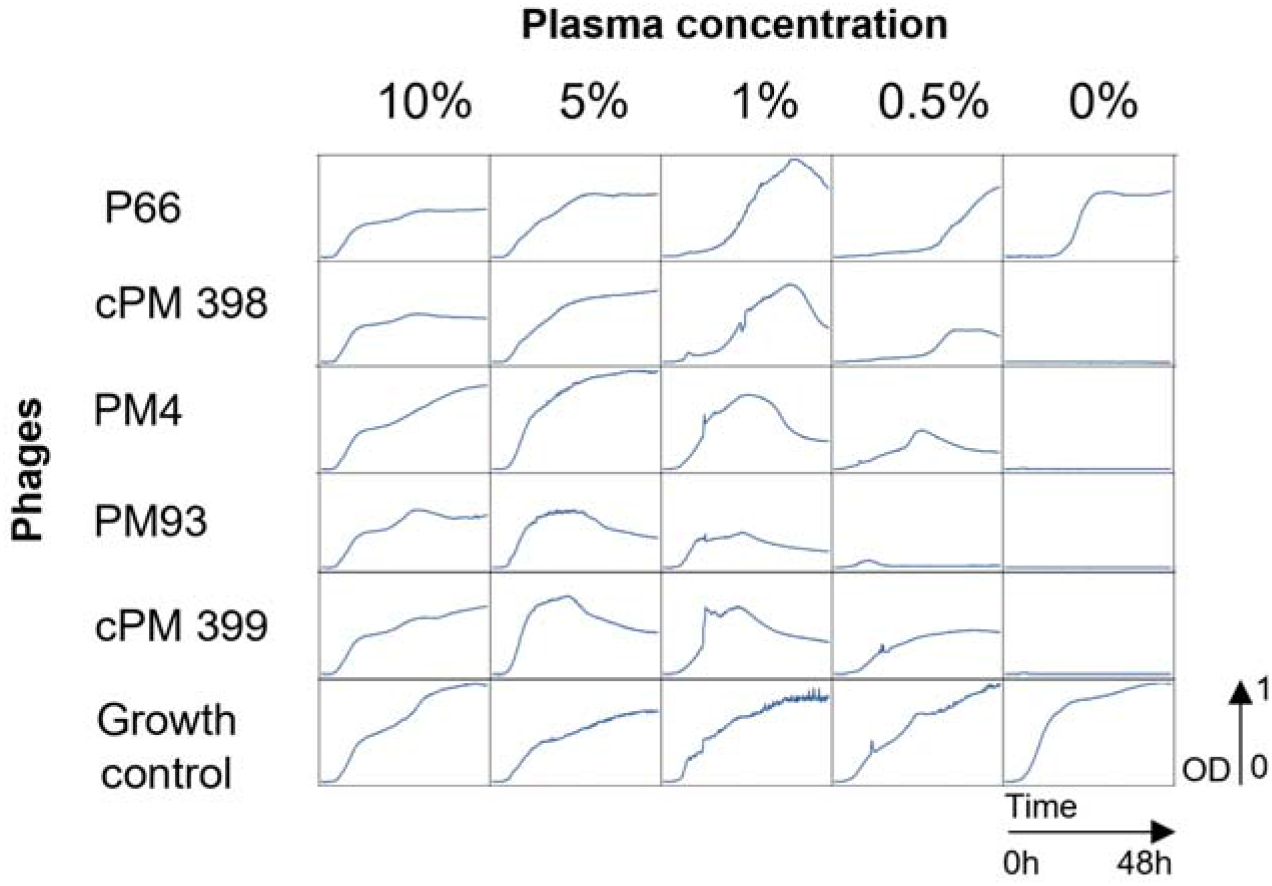
48h of growth inhibition assay in human heparin plasma (Red Cross, mix of two donors, blood groups A+/0+) The graphs show turbidity assays of bacterial suspensions at a starting bacterial concentration of 2.5×10^6^ CFU/ml in presence or absence (growth control) of the different phages or phage cocktails at 2.5×10^8^ PFU per ml (MOI = 100). The mixture was incubated for 48h at 37 °C at different concentrations of human plasma as indicated on top of the graph. CFU, colony forming units; MOI, multiplicity of infection; OD, optical density at 600nm; PFU, plaque forming units.

As observed in human plasma, activity of phage cocktails and single phages was also severely impaired in the presence of guinea pig plasma (Supplementary Fig. 1).

### 2.2. Phages are stable in plasma concentrations up to ∼50%

We therefore tested the possibility that the loss of phage activity could be due to a direct interaction of the phages with the plasma or the anticoagulant. We incubated various described *Herelleviridae* phages with increasing concentrations of heparin or citrate plasma and tested their residual activity after 24h. All phages tested showed < 1 log PFU/ml titer reduction after 24h incubation at 37°C in 50% heparin or citrate plasma (Fig. 2). At plasma concentrations > 50%, the phage titers dropped by up to 4 log PFU/ml.

**Figure 2.**
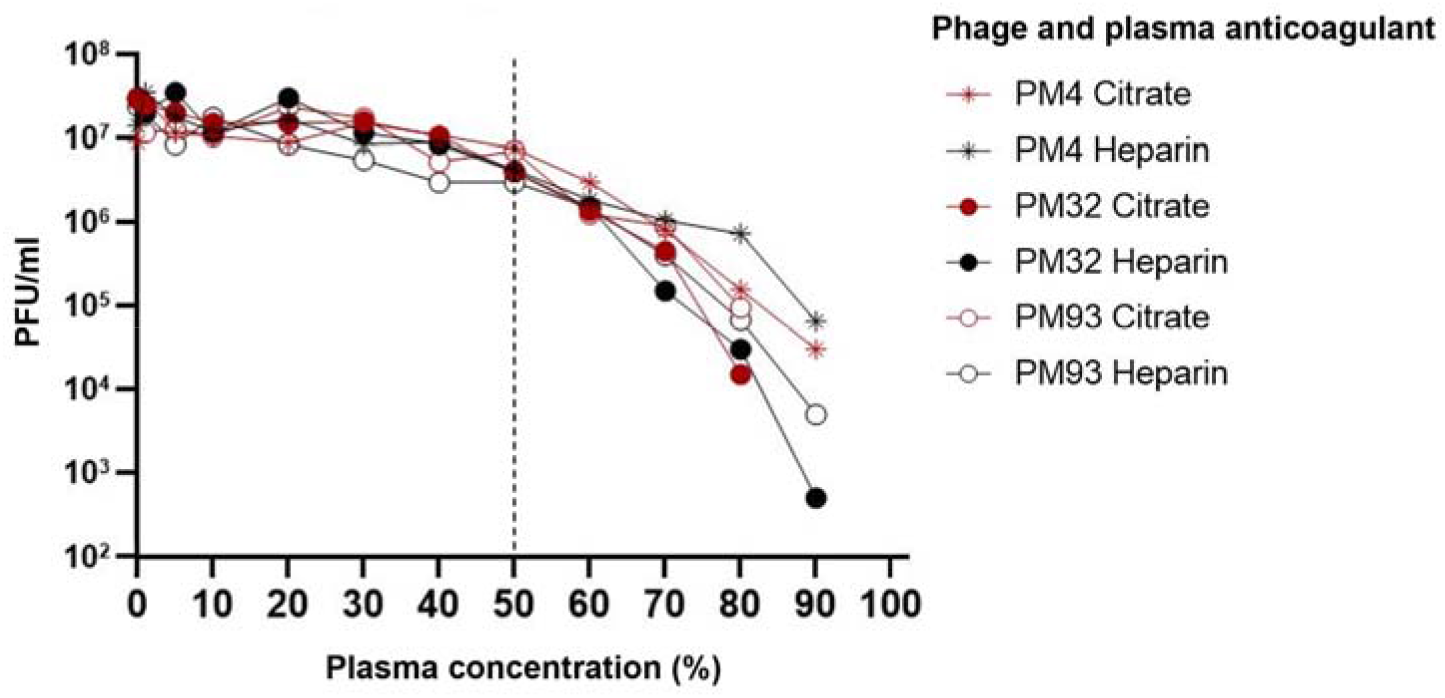
Stability of phages in human plasma. Phages were incubated for 24h at 37°C in BHI supplemented with human plasma as indicated, after which phage titer was determined by spot test on their common S. aureus host strain 124605 ^[9]^. Dotted line indicates 50% plasma concentration.

These results suggest that direct interactions between plasma or anticoagulant and phages may play a role in their instability at high plasma concentrations, but that neither plasma nor anticoagulant (data not shown) reduce phage activity at concentrations below ∼50%. Therefore, the reduced phage activity observed at a plasma concentration as low as 0.5 to 10% (Fig. 1) cannot be explained by a direct interaction between phages and plasma or anticoagulant.

### 2.3. Phage replication is not substantially impaired even though bacteria are not cleared

In phage propagation experiments, we used an input bacterial titer of 5×10^6^ CFU/ml and PM93 at an initial MOI of 0.1. We measured the phage titer after 24h incubation as a function of plasma concentration. Phage PM93 was selected since its activity was less impaired in plasma compared to the other phages (Fig. 1). Phage PM93 propagated in plasma in all the plasma concentrations tested, with a maximum increase of 4 logs at 7.5% plasma (Fig. 3a). The bacteria were able to grow despite efficient propagation of the phages in presence of ≥ 2.5% plasma (Fig. 3b).

**Figure 3.**
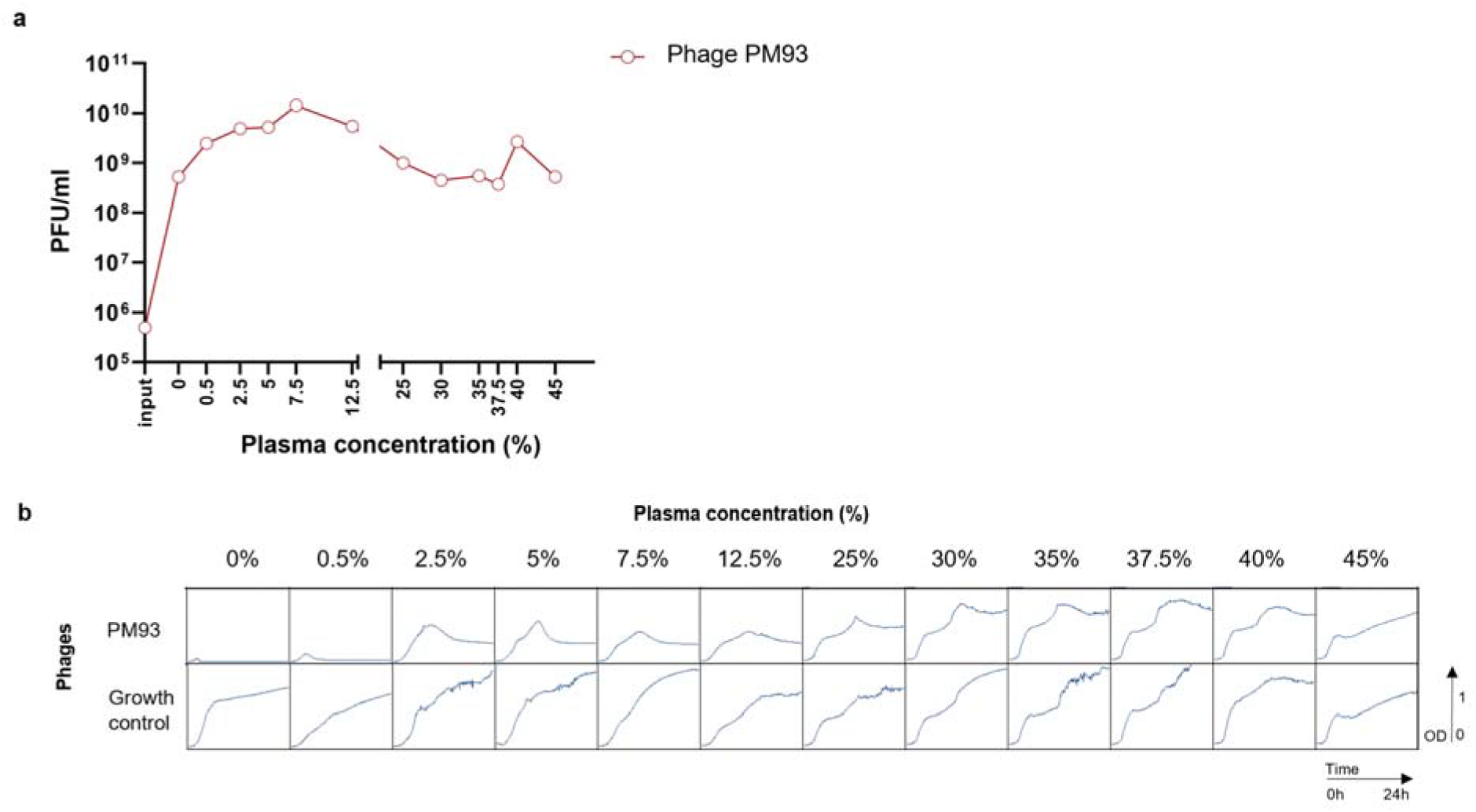
**(a) Phage propagation in plasma.** The graph shows phage titer after 24h incubation in function of plasma concentration. (b) 24h of growth inhibition assay in human plasma. The graphs show OD600 of bacterial suspensions in presence or absence (growth control) of PM93 at an MOI of 0.1. The mixture was incubated for 24h at 37 °C at different concentrations of human heparin or citrate plasma as indicated on top of the panel.

These results suggest that in plasma, although phages can still propagate, there are bacterial cells that are surviving the phage attack. We hypothesized that this may be due to a reduced adsorption rate, as suggested by previous studies ^[14]^. As a next step, we investigated whether this could be caused by masking of the surface of S. aureus by substances in plasma.

### 2.4. Phage adsorption is impaired in plasma

To test if adsorption was impaired, we measured the efficiency of the center of infection (ECOI) in presence of plasma. In ECOI, one plaque corresponds to a bacterium successfully infected with a phage ^[20]^. Due to the excess of bacteria relative to phages, ECOI is very sensitive to changes of low adsorption rates (while the classic phage adsorption assay ^[21]^ is more suitable to measure [log-scale] reductions in phage titer, indicating a high rate of adsorption).

As shown in Fig. 4, 10% plasma in BHI reduced the adsorption of the phages by 1.8-log infected CFU/ml compared to the same phages in BHI alone, suggesting that human plasma substantially affects the adsorption of the phage onto *S. aureus* cells.

**Figure 4.**
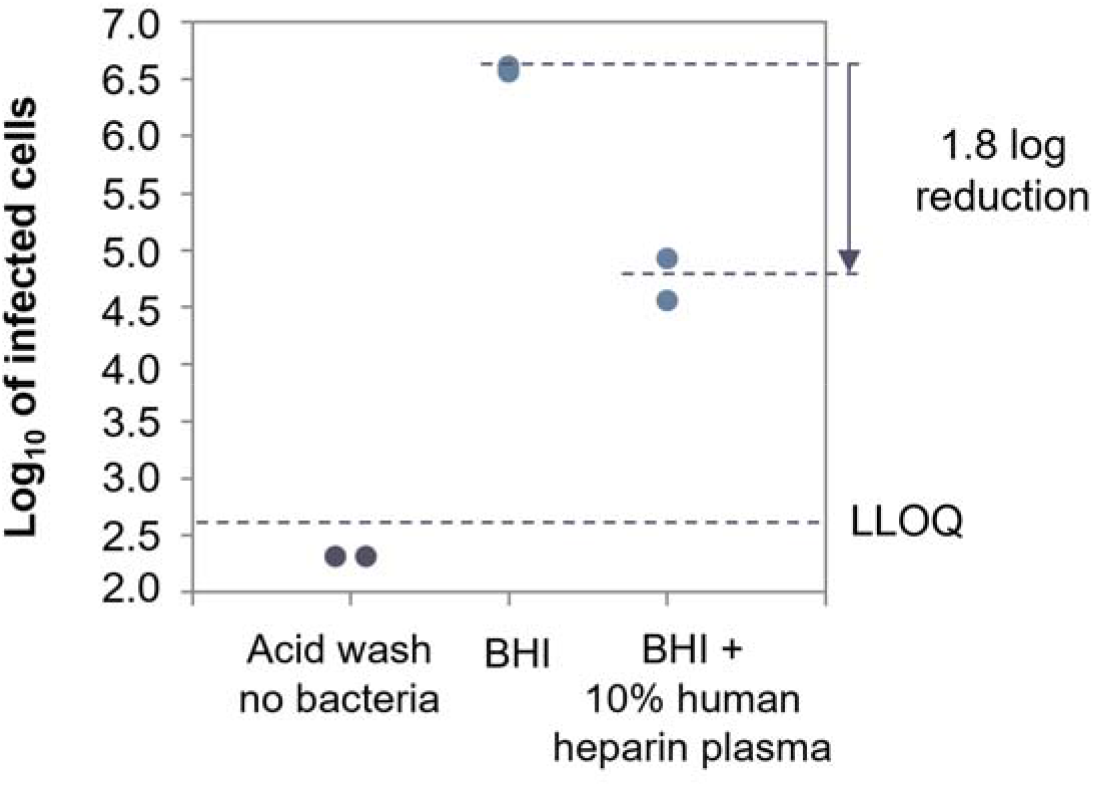
ECOI of *Herelleviridae* phage PM93, at MOI of 0.01 in human heparin plasma. The log 10 of the number of infected cells is plotted against the different conditions tested in the assay: control without bacteria, BHI, BHI+10% plasma. LLOQ: Lower limit of quantification.

### 2.5. Purified fibrinogen is sufficient to impair phage activity

Since phage adsorption was impaired in the presence of plasma, we tested if this was related to the ability of *S. aureus* to form clumps, which is induced by fibrinogen ^[16]^, and involves the formation of macroscopic biofilm-like structures. We first verified that purified fibrinogen in medium clumps *S. aureus* in absence (not shown) or presence of phages, starting at a concentration of 0.2 mg/mL (Fig. 5a). In the same conditions we also observed that the inhibition of the phage activity correlated with the concentration pf purified fibrinogen supplemented BHI, as shown in Fig. 5b. All tested phages (*Podoviridae* P66 and highly diverse *Herelleviridae* BT3, PM4 and Remus/Romulus derivatives PM56/PM93) were completely inhibited by the addition of fibrinogen, starting at 0.4-1mg/ml, which corresponds to 25% of its physiological concentration ^[22]^. These findings suggest that fibrinogen is sufficient to induce clumping of *S. aureus* and to inhibit the phage activity in plasma.

**Figure 5.**
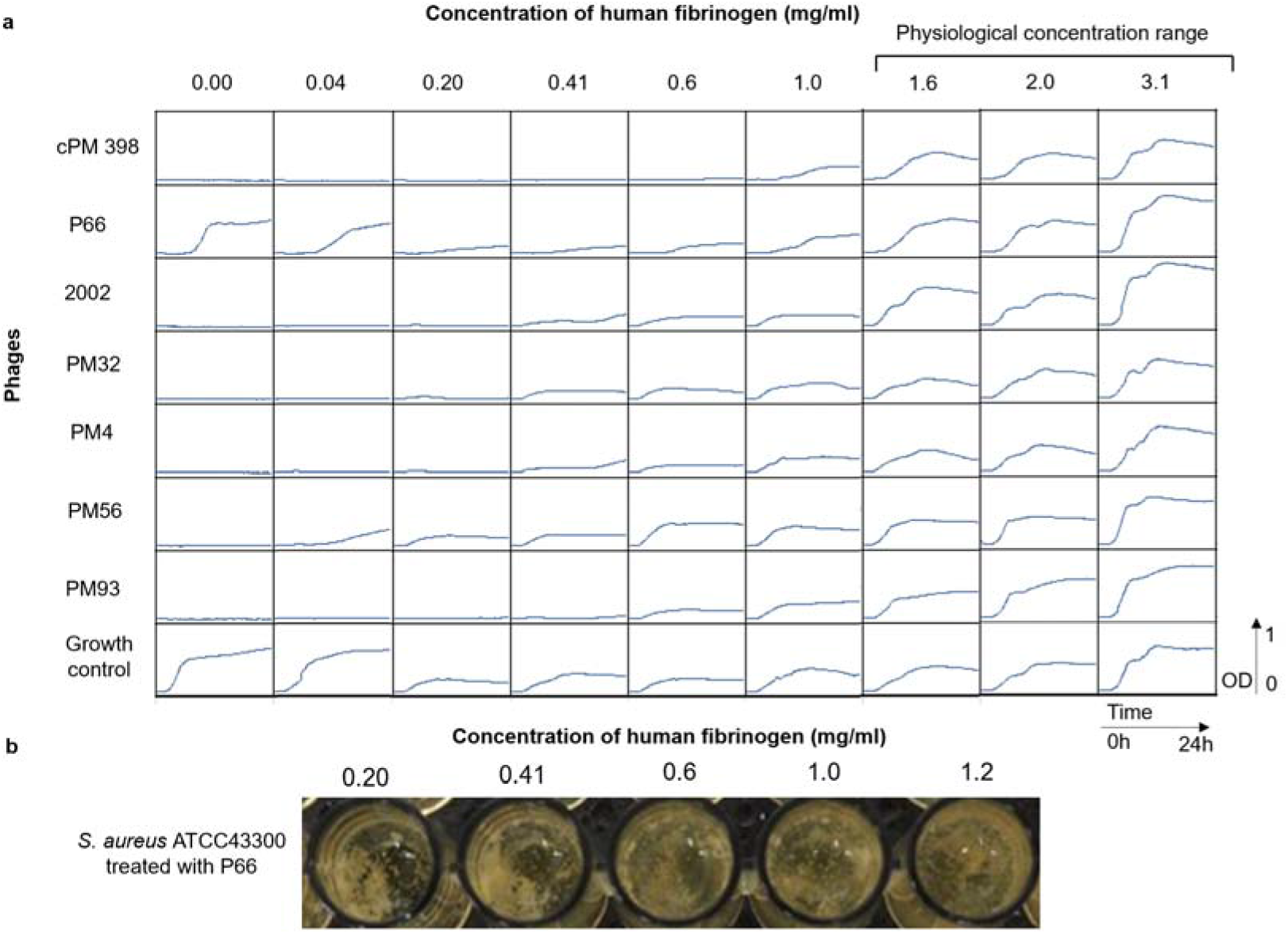
**(a) S. aureus aggregates in presence of fibrinogen**. Macroscopic clumps formed after 24h in wells treated with phage P66, with fibrinogen concentration increasing as indicated. **(b) 24h of growth inhibition assay in presence of fibrinogen**. The graphs show OD600 of bacterial suspensions in presence or absence (growth control) of the different bacteriophages as indicated on the left of the panel. The mixture was incubated in BHI for 24h at 37°C at different concentrations of fibrinogen as indicated on top of the panel. Physiological concentration range according to ^[22]^.

### 2.6. S. aureus induces clotting and tPA lyses S. aureus-induced clots in plasma of different species

Fibrinogen alone can inhibit phage activity via clumping; however, in plasma S. aureus also induces coagulation, forming local clotting around the cells, which might also hinder the phages from adsorbing to the cell surface. It was previously shown by ^[23]^ that the thrombolytic drug tPA prevented S. aureus biofilm formation in presence of plasma by blocking its attachment. We tested if fibrinolysis with tPA, might be effective to lyse the pre-formed clot and restore phage activity *in vitro*.

As shown in Supplementary Table 1 and Fig. 6, human tPA lysed S. aureus-induced clots in human heparin and citrate plasma, guinea pig citrate plasma and rabbit heparin plasma. Of note, *S. aureus* did not induce macroscopically visible clotting in murine citrate, heparin guinea pig and citrate rabbit plasma. Moreover, no macroscopic clumping was observed in murine citrate or guinea pig plasma under these experimental conditions.

**Table 1.** List of phages used in this study.

| Phage Name | Reference | Accession Number |
| --- | --- | --- |
| vB_SauH_2002 | [9] | MW528836 |
| BT3 | [9] | MW546073 |
| 812 | [33]<br>[9] | MH844528.1<br>MW546072 |
| Remus | [34]<br>[9] | JX846612<br>MW546076 |
| Romulus | [34]<br>[9] | JX846613.1<br>MW546077 |
| PM4 | [9] | MW546074 |
| PM9 | [9] | MW546064 |
| PM22 | [9] | MW546065 |
| PM25 | [9] | MW546066 |
| PM28 | [9] | MW546067 |
| PM32 | [9] | MW546070 |
| PM34 | [9] | MW546068 |
| PM36 | [9] | MW546069 |
| PM56 | [9] | MW546071 |
| PM93 | [9] | MW546075 |
| P66 | [8] | NC_007046.1 |

**Figure 6.**
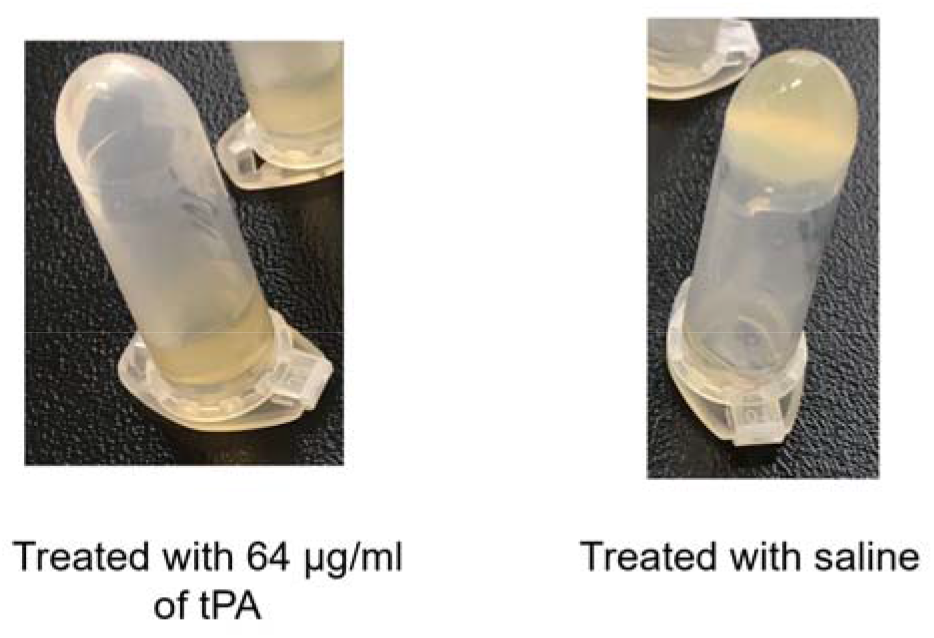
Example of tube clotting test. 400 μL of human 25% heparin plasma (diluted with saline) were clotted by adding 10 μL of S. aureus strain ATCC 43300 at OD=0.5 in suspension and incubated at 37°C overnight. The clot was then treated by adding 225 μL of 25% (end concentration) of plasma diluted with saline and 64.5 μg/mL tPA (end concentration) or no tPA (control). Where the clot was dissolved, the material collects at the bottom of the tube (clot lysed), while it sticks to the top (unlysed clot) upon treatment with saline.

### 2.7. tPA partially restores phage activity in human plasma

Since tPA was able to efficiently lyse the clot formed by *S. aureus* ATCC 43300, we tested its effect on phage activity in human plasma.

A tPA concentration of 10μg/ml was insufficient to restore phage activity at even 1% plasma (Fig. 7a). However, as shown in Fig. 7b, tPA at 64 μg/ml partially restored phage activity in up to 5% human citrate plasma.

**Figure 7.**
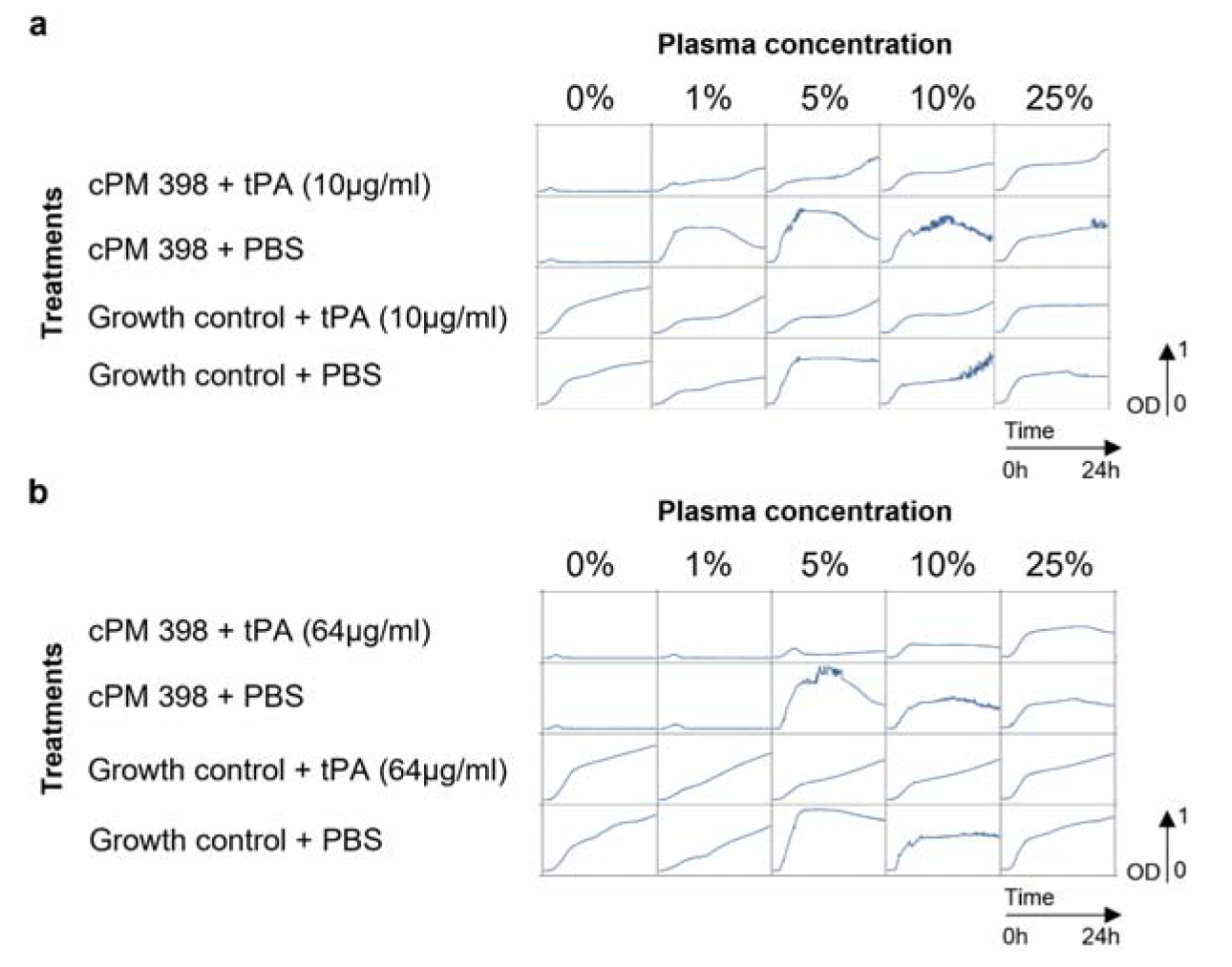
24h of growth inhibition assay of PM-398 in human citrate plasma supplemented with tPA. The graphs show optical density measurements at 600 nm (OD600) of bacterial suspensions at 5×10^6^CFU/ml in presence or absence of cPM 398 (phage cocktail of P66+ 2002, MOI 0.1) as indicated on the left and on top of the panel. The mixture was incubated for 24h at 37 °C at varying concentrations of plasma and with different concentration of tPA as indicated in the graph (a: 10 ug/ml, b: 64 ug/ml, or PBS).

Since fibrinolysis by tPA improved the phage activity, we tested whether inhibition of the coagulation, prior to clot formation, may have the same effect. Dabigatran (Pradaxa®) has inhibitory activity on the staphylococcal coagulase ^[24]^, and therefore it may prevent the clotting induced by *S. aureus*. As shown in Supplementary Fig. 2, dabigatran alone, after incubation with plasma and bacteria, was not able to improve the phage activity. This underlines the observation that aggregation of the *S. aureus* cells (e.g. by fibrinogen-induced clumping) is sufficient to block phage activity.

These results strongly indicate that clotting (a process distinct from clumping) adds too but is not required for the inactivation of phage activity in human plasma.

### 2.8. tPA completely restores phage activity in guinea pig plasma

As shown above, tPA was active on S. aureus-induced clots in guinea pig plasma, therefore we tested whether human tPA was also able to restore phage activity in guinea pig plasma.

As depicted in Fig. 8, tPA rescued phage activity in guinea pig heparin plasma, while with the highest guinea pig citrate plasma concentration there was staphylococcal growth after 12h. The stronger effect of tPA in guinea pig plasma compared to human plasma could be explained by two factors: first, the fact that it contains lower concentrations of coagulation factors ^[25]^. Second, in guinea pig plasma no clumping was observed (Supplementary Table 1).

**Figure 8.**
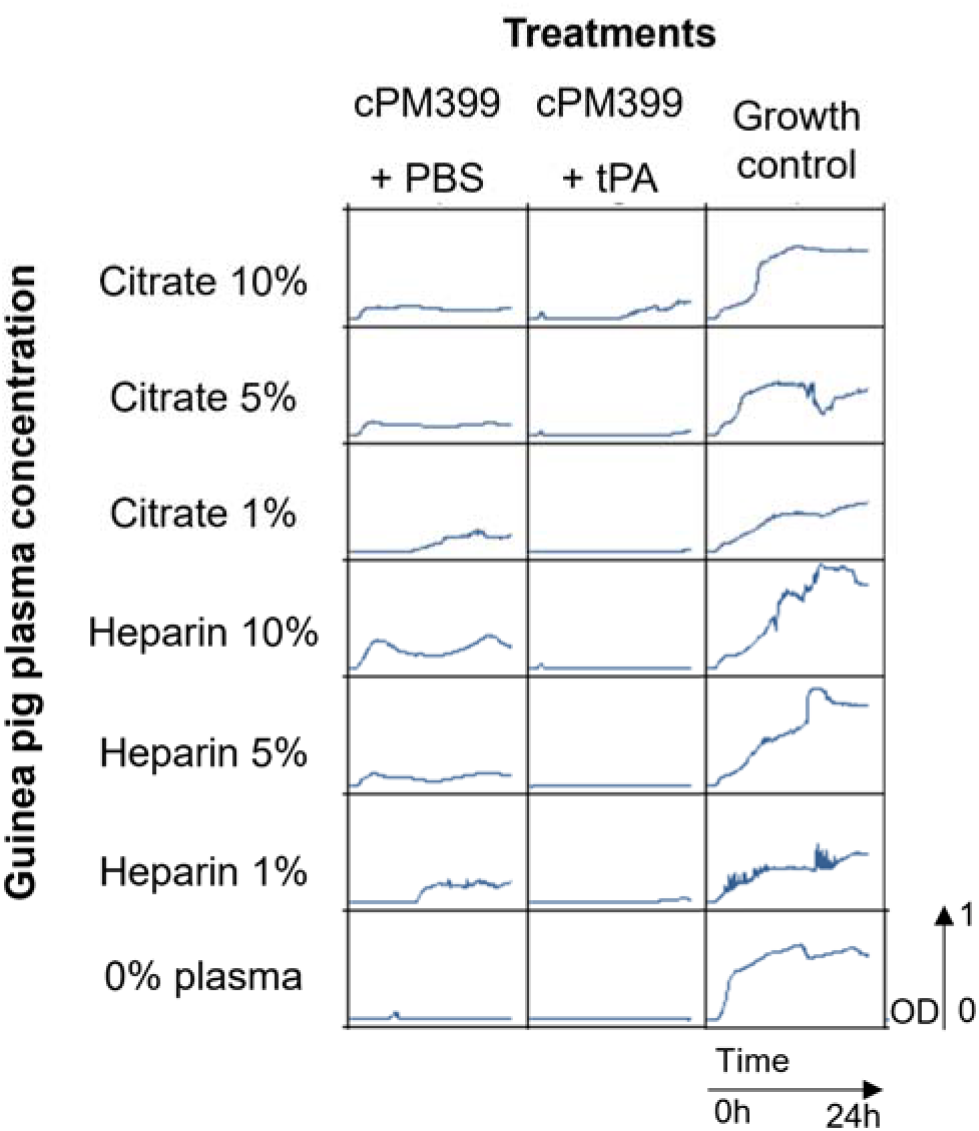
24h growth inhibition assay of PM-399 (MOI 100) in guinea pig plasma in presence or absence of tPA and plasma. The different concentrations of citrate or heparin plasma used are indicated on top and left of the panel. The graphs show optical density measurements at 600 nm (OD600) of suspensions of *S. aureus* ATCC43300 for 24h at 37 °C.

These results suggest that both clumping and clotting contribute to phage inhibition in human plasma, while in guinea pig plasma clotting (and not clumping) seems to dominate the loss of phage activity.

### 2.9. tPA improves phage adsorption in guinea pig plasma

We tested whether phage activity could be recovered by tPA via recovery of adsorption with an ECOI assay as described above. tPA increased the infectivity of phages ten times (1 log), compared to the no tPA control (Fig. 9), confirming that tPA, in guinea plasma, increases the accessibility of the phage receptor on *S. aureus* cells.

**Figure 9.**
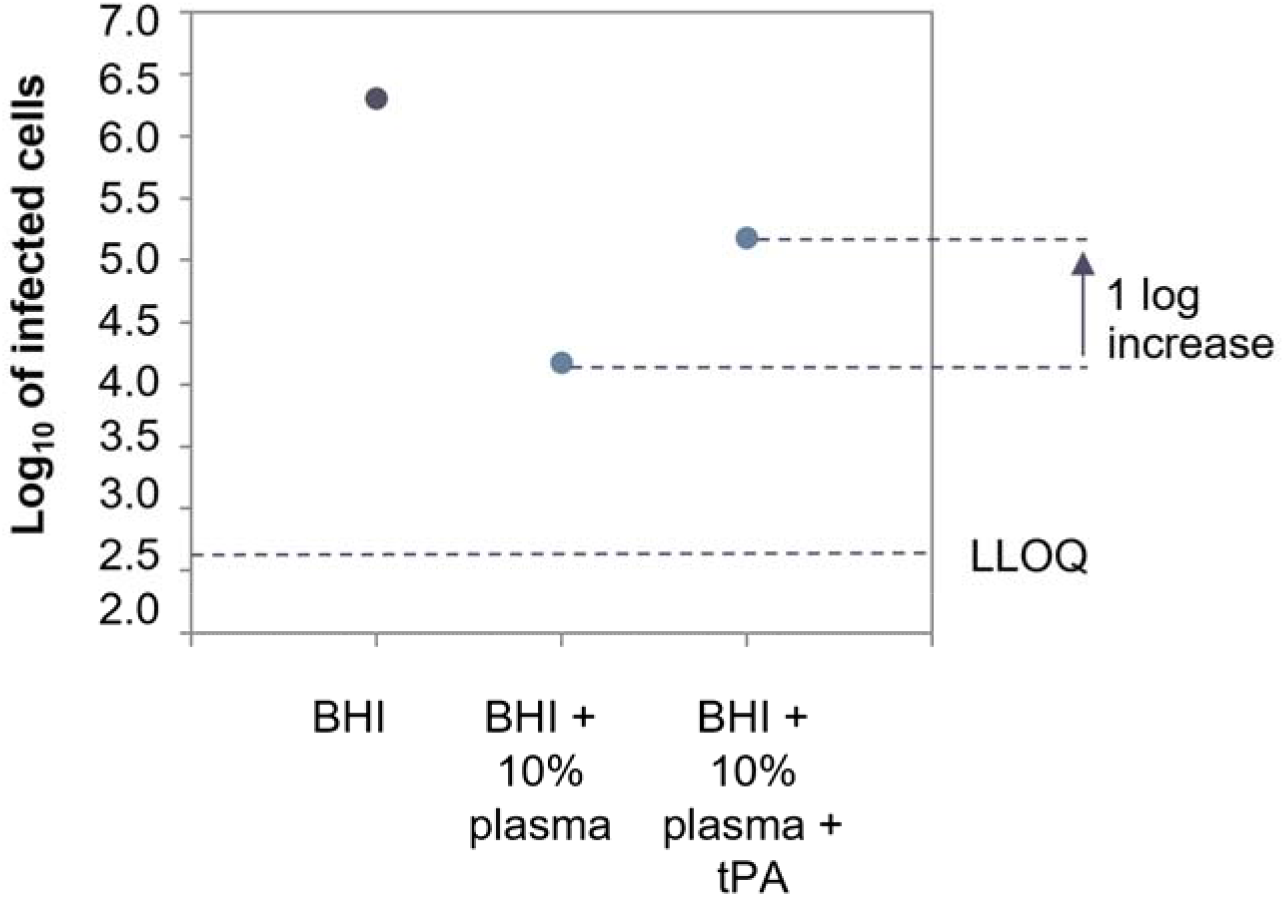
ECOI of Herelleviridae phage PM93 at MOI 0.01 in guinea pig heparin plasma. The log 10 of the number of infected cells is plotted against the different conditions tested in the assay: BHI, BHI+10% plasma and BHI+10% plasma + tPA at 64 μg/ml. LLOQ: Lower limit of quantification.

### 2.10. Phage breeding in presence of plasma does not improve the phage activity

To improve the activity of phages in plasma, a combination of phages was subjected to directed evolution (what we called breeding), by passaging them with 24 different S. aureus strains and in presence of plasma for 20 rounds as described previously ^[9],[26]^. The single phages isolated after breeding did not show a superior activity in plasma in comparison to the ancestor phages (data not shown). These results imply that to be active in plasma, the phages might need a genetic change larger than what can be achieved in this lab-scale directed evolution assay.

### 2.12. tPA restores phage activity on pre-formed S. aureus aggregates in bovine synovial fluid

In the context of prosthetic joint infections, *S. aureus* is also able to form aggregates in the presence of synovial fluid. In previous studies, ^[16]^ it was shown that these aggregates in synovial fluid led to enhanced antibiotic resistance. To revert the formation of these macroscopic, biofilm-like structures, they dissolved the aggregates with tPA at 1mg/ml (a concentration not achievable *in vivo* due to the risk of hemorrhage) thereby successfully restoring the effectiveness of antibiotics.

We tested if phage lytic activity in synovial fluid could also be restored with tPA. Macroscopic clumps were formed by incubating 3×10^7^ CFU of S. aureus ATCC43300 at 37°C for 24h in 70% bovine synovial fluid, and then treating these with a combination of tPA and either phage cocktail cPM399 or a combination of vancomycin and rifampicin at 10-fold MIC. As depicted in Fig. 10a, *S. aureus* formed large aggregates in bovine synovial fluid after 24h, consistent with the observation by ^[16]^ in human, equine and porcine synovial fluid.

As depicted in Fig. 10b, none of the single treatment alone could dissolve the clots. However, tPA treatment alone led to turbid solutions with aggregates (as compared to clear solutions without tPA), indicating a partial de-aggregation of *S. aureus* clumps, consistent with the findings of ^[16]^. The treatment with the combination of cPM-399 and tPA led to dissolution of the aggregates and a clear solution, pointing at a strong synergy between tPA and the phage cocktail cPM-399. Interestingly, the combination of tPA and antibiotics did not induce dissolution of the aggregates.

The treatment with phage cocktail (cPM-399) also led to a statistically significant reduction of bacterial viability by 1.5 log CFU/ml versus control (Fig. 10c). Consistently with the turbidity observation depicted in Fig. 10b, tPA treatment alone did not reduce the CFU/ml. In contrast, combined treatment with tPA and cPM-399 led to a median reduction of 5 log CFU/ml. This demonstrates a strong synergy between tPA and the phage cocktail cPM-399.

We also measured the phage counts (PFU/ml) after the treatment and found that, consistent with the results depicted in Fig. 10a to c, the phages replicated approximately ten times more (1 log PFU/ml) in the wells treated with the combination of tPA+cPM-399 compared to the wells treated with PM-399 only (Fig. 10d).

These results are in agreement with the findings of Gilbertie et al (2019) ^[16]^, who showed that various antibiotics were not effective in killing the bacteria as long as aggregates could be observed macroscopically.

Whether tPA can be used to enhance the efficacy of phages to treat S. aureus infections in humans largely depends on the risk of hemorrhaging. In this respect, the approach shown here indicates a pathway to reduce the effective concentration of tPA to 64 μg/ml, if combined with phages, as compared to the combination with antibiotics (tPA at 1000 μg/ml) described in ^[16]^.

### 2.14. Phages were inactive in a mouse S. aureus thigh infection model

Given that in vitro, phage activity was suppressed in presence of plasma, we evaluated whether this was relevant *in vivo*.

First, we assessed the pharmacokinetics (PK) of phage PM4 in female NMRI mice. The same dose of PM4 (2.6×10^10^ PFU) was administered via intraperitoneal (i.p.) or intravenous (i.v.) injection, followed by euthanasia after 30 min, 2h or 6h.

In the thigh, the C_max_ was 1.8×10^6^ PFU/ml after 30 min after i.p. administration of PM4, compared to a C_max_ of 8.8×10^6^ PFU/ml at the same time point after i.v. administration. The difference between administration routes was not statistically significant (Supplementary Fig. 3). In plasma, the C_max_ was 1.8×10^8^ PFU/ml 30 min after i.p. administration of PM4, compared to a C_max_ of 1.2×10^9^ PFU/ml at the same time point after i.v. administration. Also in this case, the difference between administration routes was not statistically significant.

The phage distributed effectively across the different tissues, although the titers in the thigh were lower compared to the other organs, where the titers decreased over time from 10^4^-10^7^ PFU/g after 30min, to 10^3^-10^5^ PFU/g at 6h (Supplementary Fig. 3).

Next, we assessed the efficacy of PM4 and one of its ancestor wild type phages, 812, in a thigh model in neutropenic mice (to avoid a potential synergy with the immune system), using the antibiotic vancomycin as a control. Two doses of 7.5×10^9^ PFU or 60 mg/kg vancomycin were administered at 2h and 8h after infection, respectively, and mice were sacrificed 24h after infection (16h after the second dose). As shown in Fig. 11, and despite presence of phages demonstrated in the PK study, neither PM4 nor 812 significantly reduced the bacterial burden in the thigh, compared to the control group. Vancomycin reduced the bacterial burden by about 4 log CFU/g tissue compared to vehicle treatment.

**Figure 10.**
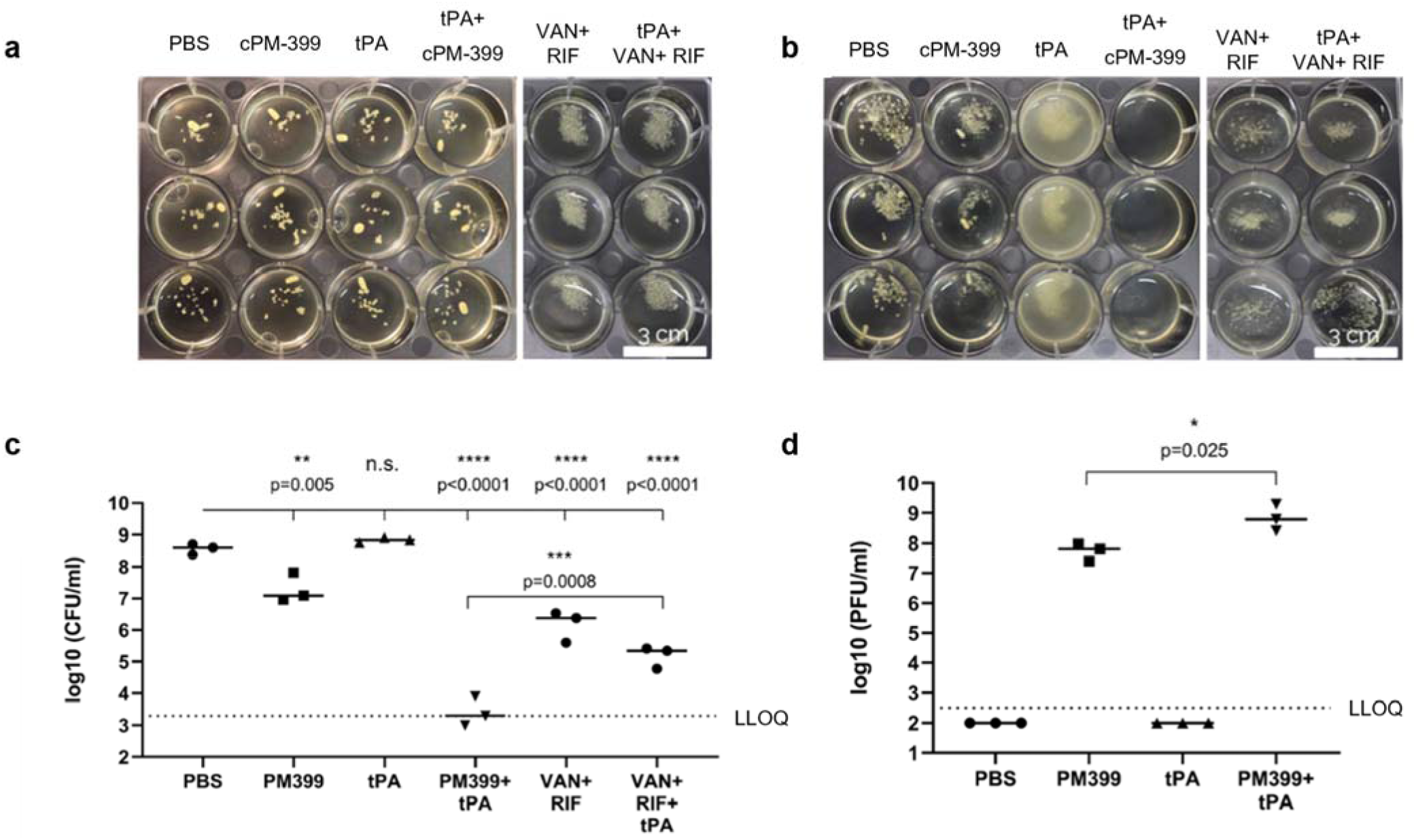
Effect of tPA in synovial fluid in combination with phages or antibiotics. (a) Clumps of *S. aureus* strain ATCC 43300 in bovine synovial fluid after 24h of incubation (in triplicate). (b) Same samples of (a) 24h after the treatment. (c) CFU/ml counts of samples in (b), p-values were calculated with a two-tailed one-way ANOVA. (d) PFU/ml counts of samples in (b), p-value was evaluated with an unpaired t-test. LLOQ: Lower limit of quantification.

**Figure 11.**
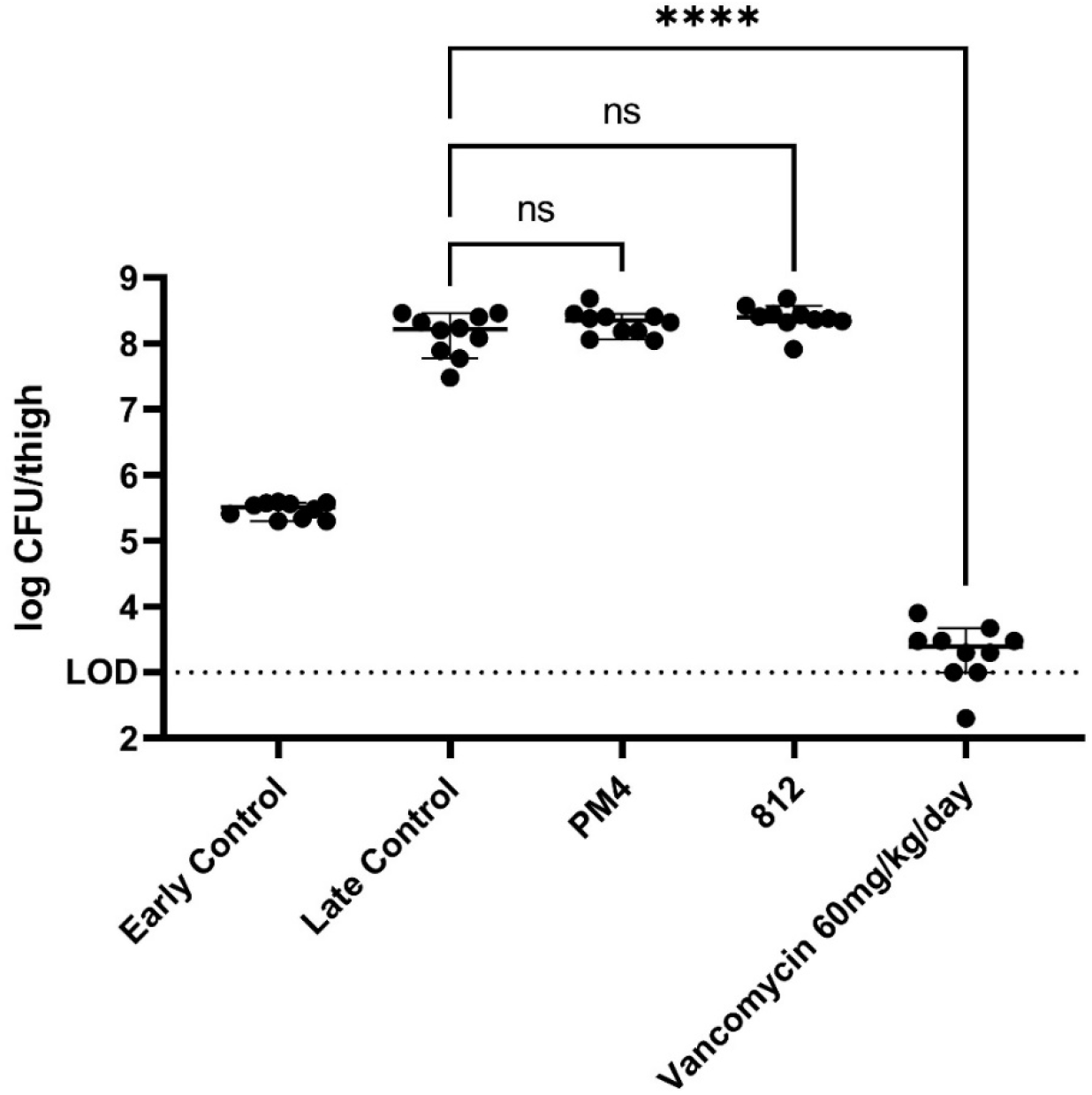
Bacterium log CFU retrieved per thigh in vivo after phage treatment per treatment group.

These results suggest that the phage-host interaction is affected also *in vivo*, most probably due to the same phenomenon observed in vitro: the phages are not able to either reach the bacteria in the infected animals, or, more likely, the bacteria are protected from the phages in the environment of the blood and interstitial tissues. This latter interpretation fits well with the data shown previously, indicating that S. aureus binds elements of the coagulation system, blocking adsorption of the phages.

## 3. Discussion

In this study, we analyzed the activity of staphylococcal phages in presence of physiological fluids (plasma and synovial fluid). We show that both plasma and synovial fluid of various mammals, including humans, impair the activity of any phage tested, of taxonomic families as diverse as *Podoviridae* and *Herelleviridae*. We thereby substantially extend the findings described by ^[12],[13]^. Although the phages could not prevent the growth of bacterial populations in presence of these physiological fluids, we found that phages were propagating under these conditions, and were stable at plasma concentrations below 50%. This suggests that the impairment of bacterial growth inhibition by the phages *in vitro* is not due to a stability issue, but rather to an altered phage-host interaction.

Shinde P. *et al*, hypothesized that this inhibition might be due to the masking effect of the coagulase activity, and/or masking by the anti-teichoic acid IgG antibodies ^[12]^. However, our data indicate that reduced phage-host absorption via a combination of clotting (e.g. *via* coagulase A) and clumping (e.g. *via* ClfA and ClfB) most likely reduces phage activity. Given the complex and interconnected mode of action of these virulence factors, we evaluated the role of the two pathways separately.

First, we tested the effect of fibrinogen-dependent clumping, driven by ClfA and ClfB, on the phage activity, by adding human purified fibrinogen to the medium. Fibrinogen-dependent clumping leads to the formation of biofilm-like aggregates, which play a pivotal role in the pathogenesis, and decrease the susceptibility of S. aureus to antibiotics ^[15]^. Purified fibrinogen added to medium was sufficient to inhibit phage activity, and to form macroscopic clumps. Interestingly, the clumping activity of *S. aureus* was observed in human plasma, but not in guinea pig or murine plasma. Despite the lack of clumping activity in the latter species, phage activity was also inhibited.

Therefore, we tested if coagulation could also play a role. We assessed this by testing both the destruction of the clot by tPA, and by preventing its formation with dabigatran. The addition of tPA to human plasma improved phage activity, particularly at low concentration of plasma. Moreover, the supplementation with tPA increased phage adsorption by one order of magnitude, supposedly promoting phage propagation and reducing growth of the bacterial population. In contrast, the addition of dabigatran did not improve phage activity in plasma. These apparently contradictory results may be explained by the fact that tPA activates plasmin, which in turn not only degrades fibrin in clots (coagulase A -dependent), but also degrades the fibrinogen that mediates clump cohesion (ClfA- and ClfB-dependent)^27],[28]^. tPA therefore induces the destruction of both clots and clumps.

This suggests that while the clumping activity in human plasma is sufficient to impair the phages, the coagulation is not, and both effects may be additive or synergistic. In guinea pig plasma, we did not observe any macroscopic clumping, suggesting that clotting plays a more dominant role in this species, and it explains why tPA is very effective in guinea pig plasma.

We also tried to breed the phages in plasma, as previously described to improve virulence and kinetics of propagation in medium ^[9],[26]^. Despite the high number of phages used and 20 rounds of passaging, we did not observe any improvement. This indicates that the genetic changes required in the phages to overcome the adsorption inhibition could not be achieved in this setup.

We demonstrate that the inhibitory effect on phages is not limited to plasma but extends to synovial fluid (which does not contain any anticoagulants). This can be explained by the presence of fibrinogen ^[15]^ and other elements of the coagulation system in this fluid ^[29],[30]^. However, the inhibition was impaired only when more than 50% synovial fluid was present, consistent with the lower concentration of coagulation proteins compared to plasma ^[29],[30]^.

To dissolve aggregates of *S. aureus* in synovial fluid, Gilbertie et al. (2019) ^[16]^ tested a dose of 1 mg/ml of tPA to restore antibiotic activity, a concentration that would incur a strong risk of hemorrhaging *in vivo* ^[31]^. Our data indicate that the synergy between tPA and phages would allow a substantial reduction in the effective concentration of tPA (from 1mg/ml to 64 ug/ml) to dissolve the macroscopic *S. aureus* clumps forming in synovial fluid and killing the bacteria.

To validate our conclusions *in vivo*, we tested phage activity in a thigh infection model in neutropenic mice preceded by a PK study to assay the bioavailability and distribution of PM4 phages administered i.v. or i.p..

Consistent with the hypothesis that phage adsorption is inhibited in physiological fluids containing elements of the coagulation system, no CFU reduction was observed between phage treatment and control, despite good bioavailability of phages in all tissues tested.

In a previous study ^[32]^, a 4-phage-cocktail including 2002 (ancestor of PM4 used in our study) was effective in reducing bacterial densities in pneumonia in rats. This suggests that in certain environments, such as the lungs, phages may not be inhibited.

Our experiments also highlight the importance of testing phage activity *in vitro* in conditions that resemble the physiological state as much as possible. Moreover, the study of staphylococcal aggregates in body fluids such as plasma and synovial fluid, along with the study of biofilms, is an imperative to further develop phage therapy for infections such as PJIs, chronic wound infections, osteomyelitis, soft tissue abscesses, and endocarditis. Clumping and clotting of bacteria in synovial fluid was observed also for bacteria other than *S. aureus*, such as, *Streptococcus zooepidemicus, Escherichia coli and Pseudomonas aeruginosa* ^*[16]*^, therefore investigations to assess the activity of phages against other bacteria in body fluids would be important.

The reduced activity of staphylococcal phages in plasma as described here might be a fundamental problem for phage therapy in humans. We therefore encourage further investigations.

## 4. Materials and Methods

### 4.1. Ethics

This research was conducted following approval from the Austrian Agency for Health and Food Safety Ltd. All studies were carried out in accordance with local regulations.

### 4.2. Bacterial Strains and bacteriophages

All bacterial assays described were performed with bacterial cultures of ATCC 43300. The cultures were prepared by inoculating the bacteria from TSA plates into BHI medium. The mixture was incubated for 4 to 6h at 37°C and 200 RPM. Prior to the start of the experiment, bacteria were diluted in BHI to reach the wanted OD600.

All bacteriophages used in this study are listed in Table 1.

### 4.3. Reagents

Plasminogen (#10874477001, Merk, Germany) was reconstituted by adding DPBS (#14190250, Thermo Scientific, UK). tPA (Actilyse® 50 mg or Actilyse® Cathflo® 2 mg, Boehringer-Ingelheim, Germany) was reconstituted by adding sterile water, to reach a final concentration of 1 mg/ml; upon complete dissolution of the cake, glycerol was added to a final concentration of 25% as cryoprotectant. Normal human serum (NHS, #H6914-20ml, Merk, Germany), and bovine citrate plasma (#P4639-10ML, Merk, Germany) were reconstituted as indicated in the instructions. Fetal bovine serum (#10082139) was purchased from Thermo Scientific (Austria). Citrate plasma from healthy donor was provided by Red Cross Austria Vienna. Heparin plasma was obtained by centrifuging heparin blood (Red Cross Austria Vienna). Alternatively, heparin and citrate plasma were obtained from a healthy in-house donor by venipuncture and centrifugation of the blood. Pooled plasma and serum from human, guinea pig and rabbit and bovine synovial fluid were purchased from Dunn Labortechnik GmbH, Germany. All the biologic fluids were sterile-filtered with 0.22 μm filters, except bovine synovial fluid, for which 0.8 μm filters were used. Biologic fluids and enzymes (plasminogen and tPA) were aliquoted and stored at - 80°C, aliquots were thawed and used only once. Dabigatran was obtained from a 75 mg Pradaxa^®^ pill (Boehringer-Ingelheim, Germany) with a procedure adapted from ^[35]^. Briefly, granules (collected aseptically by opening the pill) were dissolved overnight in 9 ml saline at 37°C and 200 RPM. The day after 1 ml of the suspension was added to 29 ml of saline and incubated at 37°C 200 RPM for about 6h. The mixture was filtered with 0.22 μm filter and stored at 4°C. The final concentration of the dabigatran stock solution was 277 °g/ml (equivalent to 441 μM). If not otherwise stated, phages were propagated on *S. aureus* strains 124605, according to SOP_LAB_009 version 1.0. Vancomycin (40.96 mg/ml, Pfizer) and rifampicin (60 mg/ml, Sanofi) aliquots were stored at -20°C.

### 4.4. Phage activity kinetics on planktonic bacteria

Phage activity was measured by reading the OD620 or OD590 every 5 min for several hours at 37°C with Tecan F NANO+ or Tecan F machines. Before each reading the plate was shaken for 5 seconds with 3 mm amplitude. In all the experiments, BRANDplates®, 96-well, pureGrade™ S, PS, transparent, flatbottom plates (#781662, BRAND) were used.

### 4.5. Determination of the efficiency of center of infection (ECOI)

Adsorption of phages was assessed by determining the efficiency of center of infection (ECOI)^[20]^. Briefly, bacteria were pre-incubated with either BHI or BHI supplemented with plasma for 30 min at 37°C, and then phages (MOI of 0.01) were added. After 5 min, free phages were removed by centrifugation and washing with BHI adjusted to pH 3. After pelleting again, cells were spotted on top agar prepared with uninfected bacteria, and incubated overnight at 37°C. The day after plaques were enumerated, and the adsorption was calculated by assuming that a plaque corresponds to a single infected cell.

### 4.6. Phage stability testing in presence of plasma

Phages were incubated in BHI supplemented with plasma for 37°C, and then they were spotted on a bacterial lawn of their host, S. aureus strain 124605, and the plate was incubated overnight at 37°C. The day after plaques were enumerated, and the relative phage concentration was calculated.

### 4.7. Phage activity on pre-formed aggregates in synovial fluid and plasma and treatment with tPA

The effect of tPA on phages in synovial fluid was assessed with a protocol similar to ^[16]^, as follows. 700 μL of bovine synovial fluid was inoculated with 300 μL of S. aureus strain ATCC 43300 at OD600=0.2 in BHI, and the mixture was incubated for 24h at 37°C with shaking at 120 RPM for the formation of clumps. Then, the supernatant was removed, and the treatment was performed by adding the following reagents in order: 700 mL of synovial fluid, 100 μL of tPA (final concentration 64 μg/ml) or DPBS, and PM-399 (in BHI, final concentration of 2*10^8^ PFU/ml) or BHI, or rifampicin/vancomycin (rifampicin 0.23 μg/ml vancomycin 33.3 μg/ml, both at a concentration 10 fold their MIC). The sample was incubated for 24h at 37°C and 120 RPM. After the treatment, clumps were thoroughly resuspended by pipetting, serially diluted and plated to assess CFU and PFU counts. To detect the CFU, the sample was diluted 1:10 in PBS (100 mM KH2PO4 and 50 mM NaCl) pH 3.0 for 1 min to inactivate the phages, and then serially diluted in 1:10 in PBS pH 7.0, and spotted on BHI agar. To detect the PFU, the sample was filtered with MultiScreenHTS-GV 0.22 μm (#MSGVS2210, Merk, Germany), diluted in PBS pH 7.0 and spotted on a lawn of *S. aureus* strain ATCC 43300.

### 4.8. Breeding of phages in plasma

Phage breeding was conducted based on the principles of the Appleman’s Protocol ^[36]^, as previously described ^[9],[26]^. Briefly, three to four ancestor phages were pooled to create the input phage mixture which was used to infect each of the subpanel of 24 *S. aureus* strains (Suppl. Table 2) in a 96-well microtiter plate. The *S. aureus* strains were previously incubated 30 minutes at 37°C, in BHI supplemented with 10% of human plasma. For the first round of breeding, single phages with similar concentrations were mixed and then serial 10-fold diluted in PBS. Starting concentrations of 2×10^7^ PFU/ml of phages were mixed with bacterial overnight cultures diluted to ∼ 1×10^8^ CFU/ml. After overnight incubation at 37°C, clear lysates and the first dilution of turbid lysates were pooled, sterile filtrated and used to infect the next round of breeding. This was repeated for up to twenty rounds. Single individual phages were isolated from the bred mixture lysate by re-streaking on host strain at least four times.

### 4.9. Pharmacokinetic study in NMRI mice and thigh model of infection

Phage PM4 and 812 were purified by tangential flow filtration (TFF) prior to administration to mice.

For the PK study, female NMRI mice (29 g) received 0.2 ml of PM4 (2,6*10^10^ PFU) via i.p. or i.v. injection, followed by euthanasia with Ketamine/Xylacin (0.1 ml narcotic mix/20 g) after 30 min, 2h or 6h. The organs (liver, lungs, thighs, kidneys, and spleen) were explanted and homogenized in saline. Blood was withdrawn by cardiac puncture and collected in heparin tubes, and plasma was prepared by centrifuging the blood for 5 min at 4500 RPM. All the organ samples were centrifuged at 20,000 g for 5 min and sterile-filtered by a 0.22 μm filter plate. In order to assess the phage tires, the samples were serially diluted in saline and 2 μL drops were spotted on a lawn of S. aureus strain 124605, the plates were incubated at 37°C overnight and plaques were enumerated the day after.

For the thigh model of infection, 25 female NMRI mice were made neutropenic by administering cyclophosphamide, at 150 mg/kg and 100 mg/kg four and one day before the bacterial challenge, respectively. The animals were then challenged by intramuscular injection of 1×10^5^ CFU/thigh of S. aureus strain B409. The animals were divided in 5 treatment groups: early control, late control, phage PM4 (7,5×10 ^9^ PFU/dose), phage 812 (7,5×10 ^9^ PFU/dose), and vancomycin (60 mg/kg/day). Treatments were provided 2hs and 8hs after bacterial challenge, a summary of treatment groups is provided in Supplementary Table 3). Phages were administered i.p. and vancomycin i.v..

Early control group were sacrificed right before the onset of treatment (time 2hs). The remaining mice were sacrificed 24hs after infection and thighs were collected for the assessment of bacterial burden. Thighs were homogenized in saline and used for determination of bacterial CFU/ml in MH agar with sheep blood. For determination of phage PFU/ml samples were centrifuged at 20,000 g for 5 min and sterile-filtered by a 0.22 μm filter plate. In order to assess the phage tires, the samples were serially diluted in saline and 2 μL drops were spotted on a lawn of S. aureus strain 124605, the plates were incubated at 37°C overnight and plaques were enumerated the day after.

### 4.10. Statistics

One- and two-way Anova analysis were performed with GraphPad Prism 9.

## Supplementary Materials

### Author Contributions

Conceptualization, David Sáez Moreno, Michele Mutti, Grégory Resch, Lorenzo Corsini; Funding acquisition, Lorenzo Corsini; Investigation, David Sáez Moreno, Michele Mutti, Marcela Restrepo-Córdoba, Zehra Visram,; Methodology, Michele Mutti and Lorenzo Corsini; Project administration, Lorenzo Corsini Writing – original draft, Michele Mutti and David Sáez Moreno; Writing – review & editing, David Sáez Moreno, Michele Mutti, Grégory Resch, Lorenzo Corsini.

### Funding

This research was funded by the Austrian Research Promotion Agency (Forschungsförderungsgessellschaft), grant number 883410.

### Institutional Review Board Statement

Not applicable.

### Informed Consent Statement

Not applicable.

### Data Availability Statement

The data presented in this study are available within the article and supplementary material.

