## Supplementary material for "Phage activity against *Staphylococcus aureus* is impaired in plasma and synovial fluid"

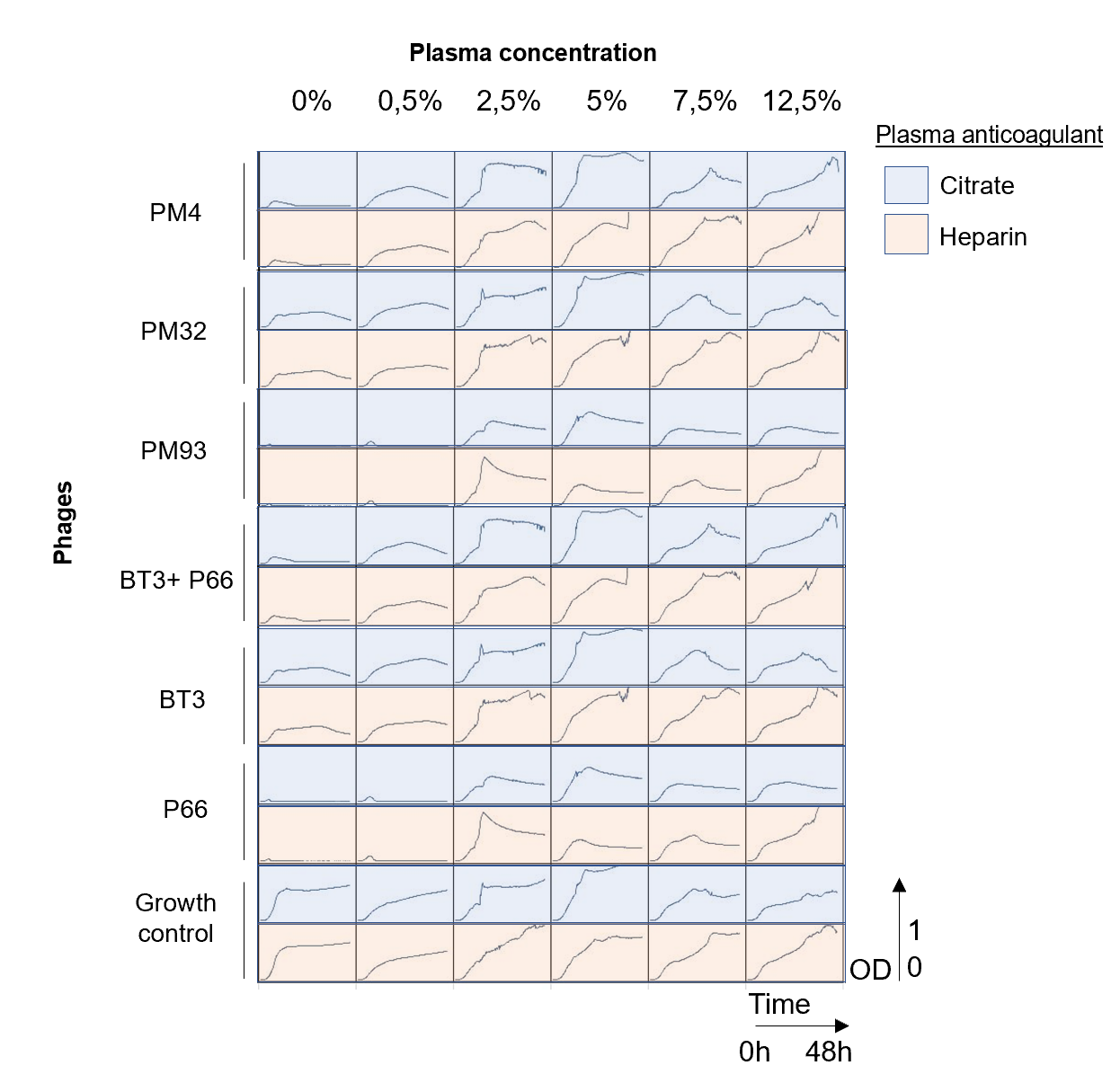

**Supp. figure 1:** 48 hours of growth inhibition assay of phage activity in guinea pig plasma. The graphs show optical density measurements at 600 nm (OD600) of bacterial suspensions of *S. aureus* ATCC43300, in presence or absence (growth control) of the different bacteriophages as indicated on the left and on top of the panel. The mixture was incubated for 48 h at 37 °C at different concentrations of guinea pig heparin or citrate plasma as indicated in the legend.

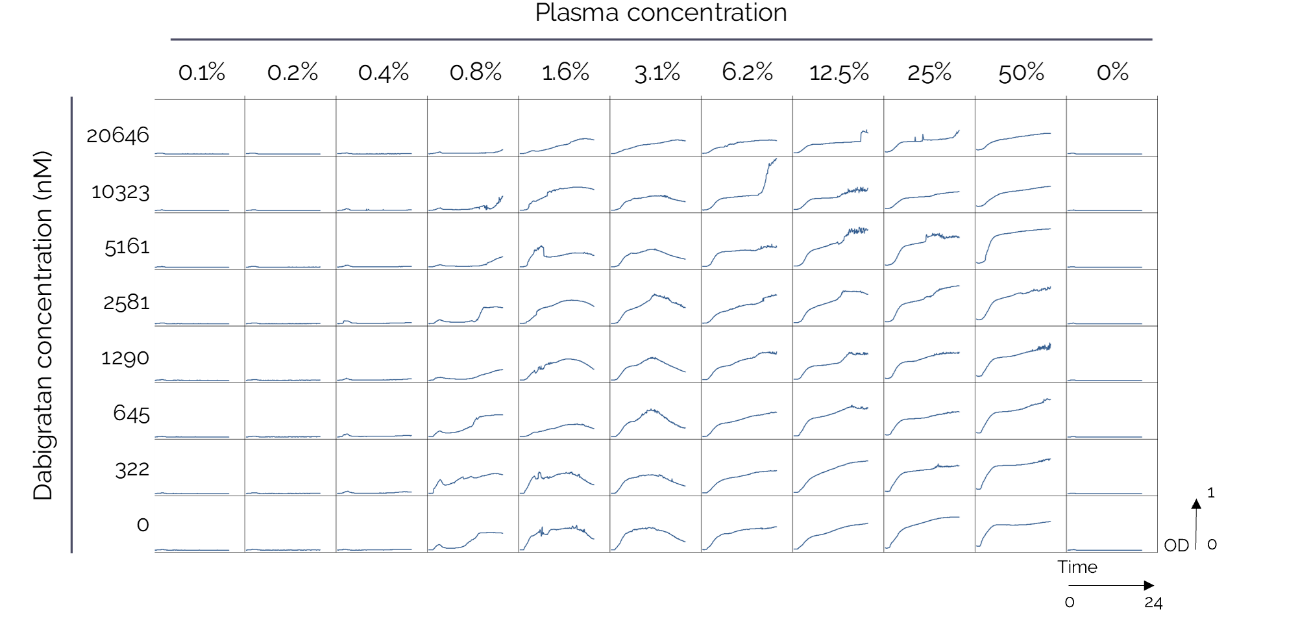

**Supplementary Figure 2.** 24 hours of growth inhibition assay of PM-398 in human citrate plasma supplemented with dabigatran (Pradaxa®). The graphs show optical density measurements at 600 nm (OD600) of bacterial suspensions in presence of bacteriophages. The mixture was incubated for 24 h at 37 °C at different concentrations of guinea pig heparin or citrate plasma and tPA, as indicated on the top of the panel . Human citrate plasma was supplemented with different amounts of *Pradaxa*® (as indicated in the legend).

**3**

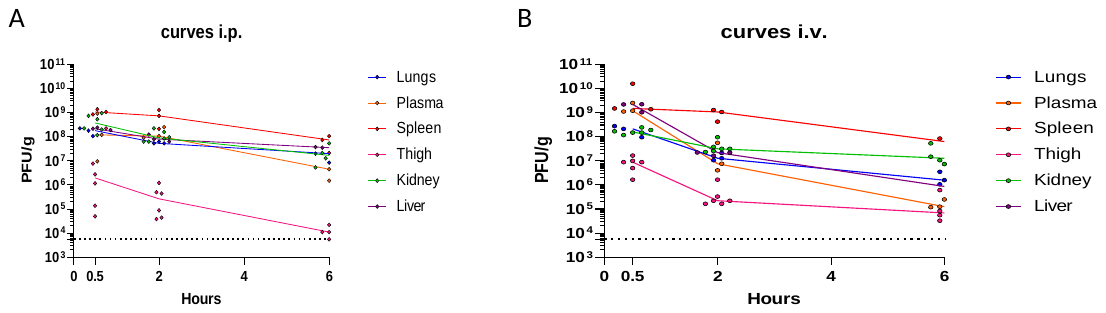

Supplementary Figure 3. A) PK of phage PM4 administered i.p. . B). PK of phage PM4 administered i.v. Each point represents a sample from a single animal, the lines connect the medians within the groups. The plasma density was assumed be 1 g/mL. To calculate CFU/g liver, one explanted lobe was weighed. To calculate CFU/g thigh, a weight of 500 mg was assumed, and 100 mg for all the other organs. The lower limit of quantification (LLOQ) indicated in the graphs refers to the thigh samples.

**Supplementary Table *1*:** Summary of the tube clot test. The tick marks indicate positive outcome, while the cross mark indicates negative outcome.

| Species | Plasma | Clumping | Clotting | Clot dissolution by tPA |
| --- | --- | --- | --- | --- |
| Human | Citrate | ✓ | ✓ | ✓ |
|  | Heparin | ✓ | ✓ | ✓ |
| Guinea pig | Citrate | 🗶 | ✓ | ✓ |
|  | Heparin | 🗶 | 🗶 | 🗶 |
| Rabbit | Citrate | ✓ | 🗶 | 🗶 |
|  | Heparin | ✓ | ✓ | ✓ |
| Mouse | Citrate | 🗶 | 🗶 | 🗶 |

**Supplementary Table 2:** *S. aureus* strains used for phage breeding.

| Clonal Complex | Name |
| --- | --- |
| CC398 | 2017-048 |
| CC398 | 2017-053 |
| CC80 | A1 |
| CC8 | A109 |
| CC15 | O111 |
| CC8 | O54 |
| CC45 | O97 |
| CC22 | A257 |
| CC30 | ATCC |
| CC398 | 2014-051 |
| CC1 | 2016-076 |
| CC22 | 2017-061 |
| CC80 | 2015-022 |
| CC22 | 2017-032 |
| CC30 | 2011-278 |
| CC45 | 2017-033 |
| CC772 | 2017-012 |
| CC5 | 2017-039 |
| CC239 | 2018-014 |
| CC49 | 124359 |
| CC88 | 124876 |
| CC45 | B42 |
| CC88 | 124896 |
| CC12 | 124882 |

**Supplementary Table 3:** Summary of the treatment groups in the tight model of infection.

*.*

| *Study group* | *Group description* | *Number of mice* | *Time of dosing 1* | *Time of dosing 2* | *Route of administration* | *Concentration* | *Volume* | *Sacrifice* |
| --- | --- | --- | --- | --- | --- | --- | --- | --- |
| *1* | *PM4* | *5* | *2 h* | *8 h* | *i.p.* | *2E+10 PFU/mL* | *0,375 mL* | *24 h* |
| *2* | *812* | *5* | *2 h* | *8 h* | *i.p.* | *2E+10 PFU/mL* | *0,375 mL* | *24 h* |
| *3* | *Vanomycin* | *5* | *2 h* | *8 h* | *s.c.* | *60 mg/kg/day* | *-* | *24 h* |
| *4* | *Early control* | *5* | *-* | *-* | *-* | *-* | *-* | *2 h* |
| *5* | *Late control* | *5* | *2 h* | *6 h* | *i.p.* | *-* | *-* | *24 h* |
